# Quadripulse Stimulation: A Replication Study with A Newly Developed Stimulator

**DOI:** 10.1101/2022.03.22.485360

**Authors:** Ikko Kimura, Yoshikazu Ugawa, Masamichi J Hayashi, Kaoru Amano

## Abstract

Quadripulse stimulation (QPS) is a promising patterned repetitive transcranial magnetic stimulation protocol, which allows the modulation of brain activity for over one hour after the stimulation. Recently, Deymed Diagnostic developed a new stimulator specifically designed to deliver QPS (DuoMAG MP-Quad, https://deymed.com/duomag-qps). The properties of the after-effect with this new stimulator were expected to parallel those obtained with another stimulator, the Magstim stimulator, which is currently used in the psychological and clinical research but not commercially available anymore. Nevertheless, experimental validation of DuoMAG MP-Quad was still warranted. We thus studied the QPS after-effect induced by this stimulator. As a result, motor evoked potentials were found to be bidirectionally modulated by the QPS for more than one hour, consistent with previous studies using the Magstim stimulator. Moreover, the degree of the after-effect was comparable to the after-effect induced by the other stimulator. Taken together, we conclude that the newly developed QPS stimulator is as effective as the Magstim stimulator.

---

Quadripulse stimulation (QPS) is a patterned repetitive transcranial magnetic stimulation (rTMS) protocol [1]. The after-effect of QPS, compared to another patterned rTMS protocol–theta burst stimulation, was reported to be stronger and less variable across participants [2]. Moreover, the after-effect of QPS is not affected by gene-polymorphisms of BDNF [3], often influencing the rTMS after-effect [4]. Therefore, QPS emerges as a promising rTMS protocol for neuro-modulational research and treatment.

Recently, Deymed Diagnostic has developed a new stimulator specifically designed for QPS (DuoMAG MP-Quad, https://deymed.com/duomag-qps), which appears as a promising QPS-stimulator [5]. However, the after-effect induced by the DuoMAG MP-Quad has not yet been systematically investigated. The degree of aftereffect may differ between QPS-stimulators considering that the waveform of stimulation pulses slightly differs between manufacturers [6]. We thus studied the QPS after-effects induced by the DuoMAG MP-Quad stimulator.

The participants were thirteen healthy adult volunteers comprising five females, 20–24 years of age (mean ± standard deviation: 21.5 ± 1.3). They were righthanded without a history of neurological or psychiatric disorders. Our study comprised two sessions with a within-subject design, applying two different types of QPS: QPS with an interpulse-interval of 5 ms (QPS5) and 50 ms (QPS50). These two protocols are expected to potentiate and depress the stimulated region most effectively [1]. The two sessions were separated by over a week, while their order was randomly assigned and counterbalanced across the participants. QPS was delivered over the left primary motor cortex (M1) using DuoMAG MP-Quad (Deymed Diagnostic s.r.o., Hronov, Czech Republic) with a butterfly-shaped 70-mm air-cooling coil (DuoMAG 70BF Air Cooled Coil; Deymed Diagnostic s.r.o., Hronov, Czech Republic). Throughout the session, motor evoked potentials (MEPs) were recorded from the right first dorsal interosseous (FDI) muscle using a Brainsight surface-electromyogram (Rogue Research Inc., Montreal, Canada).

In each session, we first localized the hotspot, defined as the location evoking the largest MEPs, for the right FDI, and measured active motor threshold (AMT), defined as the intensity evoking MEPs larger than 100 μV in 5 out of 10 trials, with a slight contraction of the right FDI (~10 % of maximum voluntary contraction). We then measured 30 MEPs at the intensity evoking ~500 μV of MEPs in the relaxed FDI as a baseline (pre-QPS). We further assessed alertness of the participants with a Stanford Sleepiness Scale (SSS; one participant presenting SSS > 4 was excluded) [7]. Subsequently, we applied QPS over the hotspot with one burst of four monophasic pulses delivered every 5 s for 30 min with a intensity of 90% AMT [1]. After applying QPS, we measured 20 MEPs every 10 min until 90 min after QPS (post-10 to post-90) using the same intensity as for the pre-QPS. The coil position was monitored with a Brainsight (Rogue Research Inc.) to confirm stable coil position.

To evaluate the overall after-effect in QPS5 and QPS50, we first compared the MEPs between pre- and post-QPS (post-10 to post-90) for each QPS protocol. The grand mean MEP amplitudes of post-QPS normalized to pre-QPS (normalized MEP) was 1.71 ± 0.41 (from below mean ± standard deviation) in QPS5 and 0.82 ± 0.23 in QPS50 (Fig. 1A). Two-tailed paired Student’s t-tests of the absolute MEP amplitude between pre- and post-QPS revealed significant enlargement after QPS5 (*P* < 0.001, *d*’ = 1.40) and reduction after QPS50 (*P* = 0.019, *d*’ = −0.80).

**Figure 1.**
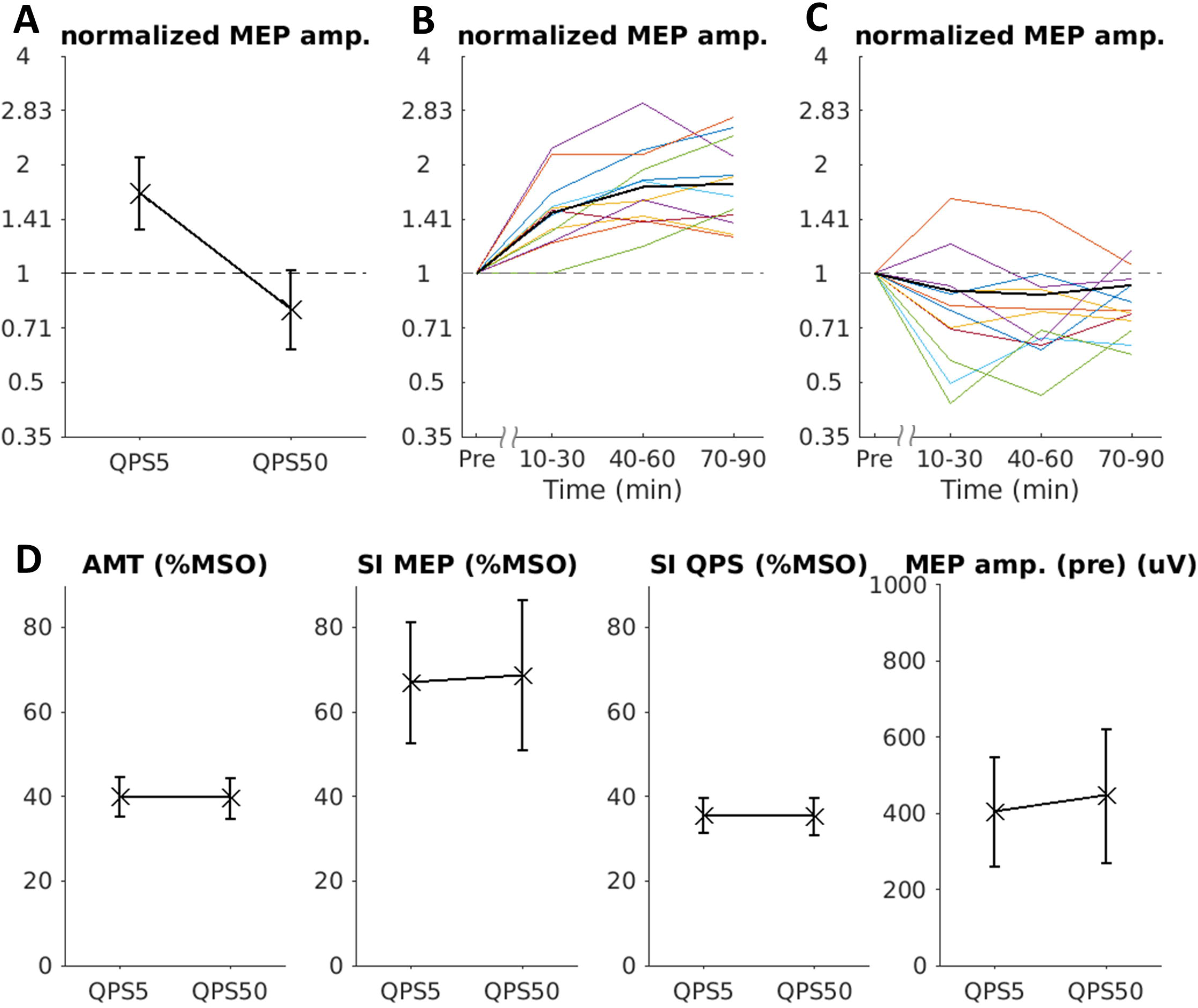
(A–C) Motor evoked potentials (MEPs) measured after QPS. (A) shows the grand average of the normalized MEP amplitudes, while (B) and (C) show the time courses of MEPs after QPS5 and QPS50, respectively. The vertical axis indicates the MEP amplitudes normalized by the baseline MEP amplitude in log scale, while the horizontal axes indicate QPS condition (A) or the time-block after QPS (B, C). (D) Comparison of baseline neurophysiological features between the QPS5 and QPS50 protocols. The active motor threshold (AMT) and stimulus intensity (SI) for recording MEPs, SI for recording QPS, and mean baseline MEP amplitude are shown. The vertical axis shows the percentage of maximum stimulator output (% MSO) in AMT and SI, while the vertical axis shows the mean baseline MEP amplitude (micro volt). The mean (black cross) and standard error (error bar) are also provided.

To evaluate the time-course of MEPs, we separated the time-course into three time-blocks (post-10–30, post-40–60, and post-70–90), then averaged MEPs among each time-block [2]. The mean of normalized MEP in post-10–30, post-40–60, and post-70–90 was 1.51 ± 0.36, 1.79 ± 0.48, 1.79 ± 0.48, respectively, for QPS5 (Fig. 1B), and 0.84 ± 0.32, 0.80 ± 0.26, and 0.83 ± 0.17, respectively, for QPS50 (Fig. 1C) (see Supplementary Fig. 1 for the individual time courses of MEPs). Repeated-measures one-way ANOVA revealed significant effect of time-block in QPS5 (*P* < 0.001, *η^2^* = 0.60) and moderate effect of time-block in QPS50 (*P* = 0.051, *η^2^* = 0.24). Post-hoc paired t-tests of the absolute MEP amplitude with correction for multiple comparisons using the Holm’s method also showed significant enlargement by QPS5 in all time-blocks (post-10–30, *P* = 0.002, *d*’ = 1.14; post-40–60, *P* = 0.001, *d*’ = 1.35; post-70–90, *P* = 0.001, *d*’ = 1.42), together with a significant reduction by QPS50 in post-40–60 and post-70–90, and a moderate reduction by QPS50 in post-10–30 (post-10–30, *P* = 0.093, *d*’ = –0.53; post-40–60, *P* = 0.030, *d*’ = –0.90; post-70–90, *P* = 0.047, *d*’ = –0.76). In terms of baseline physiological parameters, no significant differences were found in AMT (*P* = 0.73), stimulus intensity for recording MEPs (*P* = 0.49), stimulus intensity for recording QPS (*P* = 0.68) or mean baseline MEP amplitude (*P* = 0.29) (Fig. 1D) between the QPS5 and QPS50 sessions. These findings confirm steady experimental conditions across the sessions.

As expected from previous studies using the Magstim stimulator (The Magstim Co. Ltd., Whitland, UK) [1,2,8], MEPs were bidirectionally modulated for over 60 min by QPS when using the newly developed stimulator. In previous studies, the grand mean normalized MEP was ~2.0 [1], 1.60 [8], and ~1.2 [2] in QPS5, *versus* ~0.5 [1], 0.67 [8], and ~0.8 [2] in QPS50. The grand means of normalized MEP measured in the present study were comparable to those induced by the Magstim stimulator. These effects induced by the newly developed stimulator are supported by previous observations showing that short-interval intracortical inhibition is not sensitive to slight differences in TMS pulse waveform [6]. In the present study, the effect sizes of MEP-changes after QPS50 were found to be smaller than those measured after QPS5. A similar trend was reported in previous studies [1,8,9].

Together, these findings reveal that the newly developed QPS stimulator is as effective as the original QPS stimulator [1]. A promising approach to boost the aftereffect of QPS50 would be to combine QPS with afferent electric stimulation of the peripheral nerves using paired-associative QPS [10].

---

**Supplementary Figure 1.** Individual time courses of normalized MEP amplitude after QPS5 (A) and QPS50 (B). The vertical axis indicates the MEP amplitude normalized by the baseline MEP amplitude in log scale, while the horizontal axis indicates the time after QPS in minutes.

## Notes

*Declaration of interest*, None

### Competing Interest Statement

The authors have declared no competing interest.

### Summary of Updates

Co-Author's name updated

