## Supplementary Figure1 for "Quadripulse Stimulation: A Replication Study with A Newly Developed Stimulator"

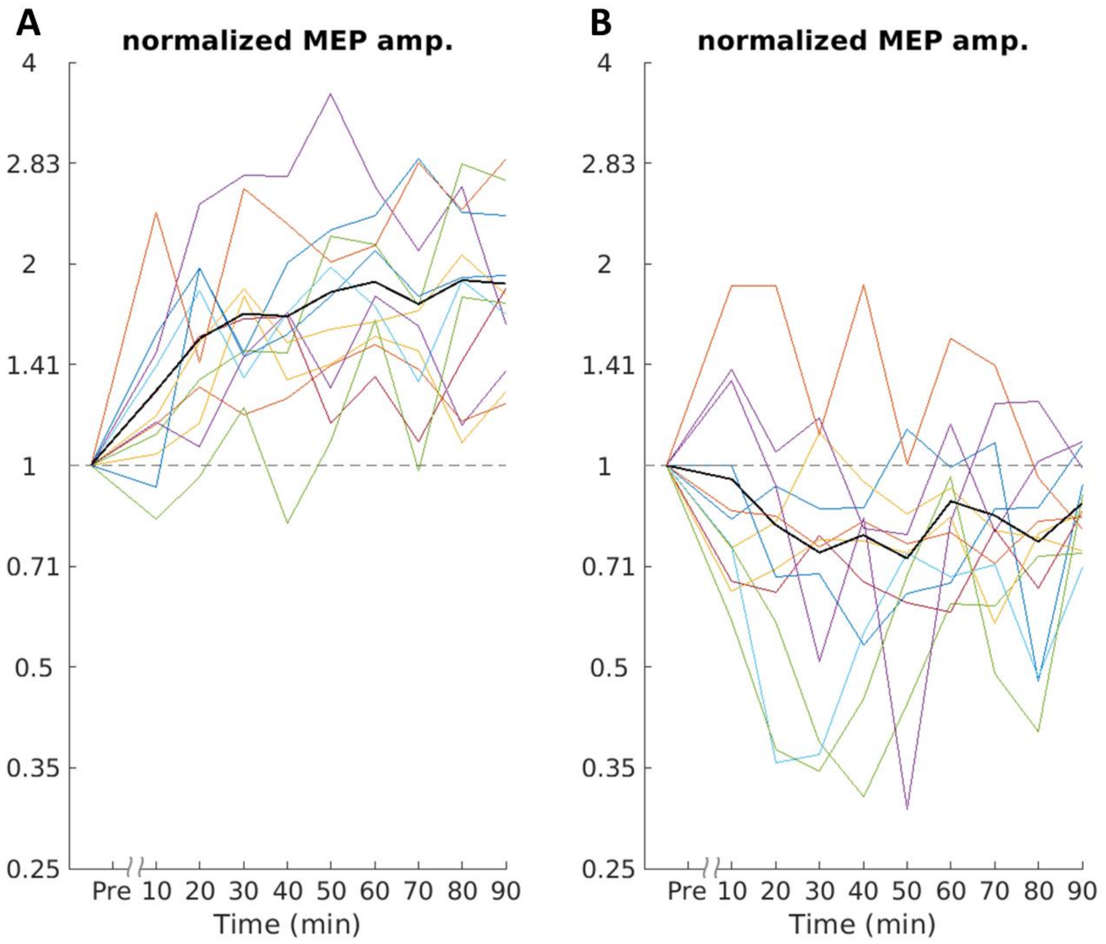


**Supplementary Figure 1.** Individual time courses of normalized MEP amplitude after QPS5 (A) and QPS50 (B). The vertical axis indicates the MEP amplitude normalized by the baseline MEP amplitude in log scale, while the horizontal axis indicates the time after QPS in minutes.
